# TILLCANN: A TILLING platform in *Cannabis sativa* for mutation discovery and crop improvement

**DOI:** 10.1101/2024.10.28.620663

**Authors:** Diana Duarte-Delgado, Konstantinos G. Alexiou, Marta Pujol, Cristobal Uauy, Nikolai M. Adamski, Victoria Vidal, Anthony Torres, Christopher Zalewski, Reginald Gaudino, Amparo Monfort, Jason Argyris

## Abstract

Cultivation of *Cannabis sativa* is increasing because of its therapeutic value and recognition as a multi-purpose and sustainable crop. Targeting Induced Local Lesions in Genomes (TILLING) is a versatile reverse genetics approach that unlocks induced variation through mutagenesis to accelerate the development of new cultivars and contribute to the functional validation of genes. Increasingly efficient next-generation sequencing technologies and genomic resources have combined to make TILLING by sequencing (TbyS) an attractive technique that can be applied in cannabis. Here we describe the development of a mutagenesis protocol for *C. sativa* and the development of the TILLCANN platform composed of 1,633 M2 families. As a demonstration of the functionality of the platform, we used TbyS to perform a high throughput screening of novel mutations in amplicons for genes associated with important agronomic and biochemical traits in a set of 512 M2 families. We confirmed 14 of the identified mutations and calculated an average mutation frequency range of 1/263 to 1/320 kb. We found that heterozygous mutants in the cannabis homologue of Class II *TEOSINTE BRANCHED 1/CYCLOIDEA/PCF* (*TCP4*) are linked to alterations in leaf number and morphology. We expect that the novel genetic variability unlocked in the TILLCANN platform for performing forward or reverse genetic screens can significantly boost breeding programs geared toward both medicinal cannabis and industrial hemp.

**Gene & accession numbers:** Raw sequencing data generated in this study was deposited at the European Nucleotide Archive with accession numbers xxx and xxx. Genes and accession numbers discussed within this manuscript correspond to the *Cannabis sativa* cs10 genome assembly deposited in National Center for Biotechnology Information with accession number GCA_900626175.2.

## Introduction

*Cannabis sativa L*., with chromosome number 2n = 20, genome size ∼ 818 megabase pairs (Mbp) for female plants and ∼ 843 Mbp for male plants (Moliterni et al. 2004) is a predominantly dioecious species distributed worldwide in a broad range of environments. Cannabis is an emblematic multi-purpose crop species as a source of fibers from stalks, oil from seeds, and phytochemicals for medicinal and psychoactive purposes mainly produced in the female flowers (Andre et al., 2016). Despite the large genetic variability present in the genus due to allogamy and cross-pollination (Salentijn et al. 2015), breeding programs have witnessed a progressive reduction of genetic variation both in medicinal cannabis and industrial hemp because of a focus on breeding for monoecious varieties and clonal micropropagation (Welling et al. 2016). Years of prohibition exacerbated this problem by preventing the establishment and preservation of cannabis germplasm in public repositories resulting in underutilization of the extant genetic variability in cultivar development (Torkamaneh and Jones, 2021).

Chemical mutagenesis employs chemical compounds that alter DNA structure, typically ethyl methanesulfonate (EMS), to produce a high frequency of randomly distributed G/C to A/T point mutations (i.e. canonical mutations). TILLING (Targeting Induced Local Lesions in Genomes) is an attractive strategy for creating novel artificial variation or diversification of traits for plant breeding that includes the induction of mutations and the screening of genes of interest to identify mutant genotypes in a large population (Tadele, 2016). TILLING is relatively simple as it does not require complicated manipulations or expensive equipment (Salgotra and Stewart, 2020). Despite its extensive use for more than 20 years in a variety of crops (Szurman-Zubrzycka et al. 2023), in cannabis only one previous study has been reported where TILLING was used in the industrial hemp variety Finola to detect mutants in fatty acid (FA) composition through PCR amplification of gene targets, heteroduplex detection, and capillary electrophoresis (Bielecka et al. 2014). More efficient, cost-effective and highly sensitive detection of mutations is enabled by TILLING by sequencing (TbyS) which couples multi-dimensional pooling of DNA with next-generation sequencing (NGS) of PCR amplicons to rapidly discover mutations present at a very low frequency in a high-throughput manner (Tsai et al., 2011). Simultaneously with detection, the nature of any induced mutation can be inferred and prioritized for analysis according to its predicted effect on gene or protein function.

Because of the change in the legal status and perception of cannabis in many parts of the world, the crop has begun to benefit from the development of important genomic resources including a high-quality reference genome from cultivar CBDRx (cs10) (Grassa et al., 2021), a recently released pangenome containing 36 haplotype-resolved, chromosome-scale assemblies (Lynch et al., 2024) and other tools (Hurgobin et al., 2021). These resources have enabled the identification of relevant homologs of target genes controlling key traits in cannabis for resistance to powdery mildew (*mildew Resistance Locus O* (*MLO*) (Stack et al., 2024; Pépin et al. 2021); candidate genes associated with fiber quality involved in the metabolism of monosaccharides, polysaccharides glycoproteins, and lignin biosynthesis (Petit et al. 2020); improved seed oil quality (Bielecka et al. 2014); photoperiod insensitivity and flowering time (Dowling et al. 2023; Leckie et al. 2023; Toth et al. 2022; Ren et al., 2021); genes associated with agronomic and morphological traits (de Ronne et al., 2024; Woods et al., 2021); and genes bearing strong selection signatures inhibiting or promoting branching and associated with variance in total cannabinoid content (Ren et al., 2021). Numerous studies have also detailed the roles of specific cannabinoid and terpenes synthases in phytocannabinoid and terpene accumulation and diversity (Booth et al., 2020; Welling et al. 2020; Allen et al., 2019; Laverty et al., 2019).

Stable transformation protocols are foundational for gene editing using clustered regularly interspaced short palindromic repeats (CRISPR-CAS9) to modify or enhance important agronomic or biochemical traits to produce improved germplasm. With a few exceptions (Galan-Avila et al., 2021; Zhang et al., 2021) cannabis has proven to be a species recalcitrant to transformation and regeneration. A lack of reliable and easily reproducible protocols for these processes means that gene editing remains an elusive goal. As cannabis is a multi-use crop with a diverse and wide-ranging set of breeding objectives, TbyS could represent a valuable strategy to overcome these constraints and to identify mutant genotypes to increase the amount of genetic variability available in breeding programs (Tadele, 2016). TbyS also provides an opportunity for functional validation of genes of interest for key agronomic and biochemical traits. Importantly, this strategy can be implemented without the biosafety concerns and policy restrictions that are relevant for transgenic and gene edited crops (Callaway, 2018; Tadele, 2016). The use of chemical mutagens and the development of TILLING platforms can therefore take advantage of more independence in terms of regulatory burdens and remain as important resources in plant breeding and genetics (Bhattacharya et al., 2023; Lucht et al; 2015).

There is no standardized and systematic protocol published for performing mutagenesis on *C. sativa* seeds, and high-throughput TbyS approaches have not previously been applied in the crop. To address these needs, we developed a cannabis EMS-mutagenized population comprised of 1,633 M2 families. This population was the foundation for the cannabis TILLING (TILLCANN) platform, employing TbyS to easily identify induced mutations in genes of interest through tridimensional pooling, amplicon sequencing, and a robust and accurate variant calling pipeline. We describe the identification and validation of mutations in genes with putative roles in powdery mildew response, flowering time, cannabinoid biosynthesis, and trichome formation in a set of 512 M2 families. Induced mutations for three of these genes were converted to allele-specific PCR markers to demonstrate their practicality for use in breeding programs. We demonstrate linkage of a mutation in a Class II *TEOSINTE BRANCHED 1/CYCLOIDEA/PCF* transcription factor, *TCP4*, to altered leaflet number and morphology, suggesting an important role for this gene in the regulation of these traits in cannabis. The TILLCANN platform will be a valuable resource to both breeders and the cannabis research community.

## Results

### Selection of *C. sativa* line and optimization of EMS application

From three F3 lines derived from the cross of a CBD-accumulating genotype and the hemp cultivar Finola, *C. sativa* line TILL8 showed the highest rates of germination and normal seedling development and was thus selected for optimizing experimental conditions for subsequent mutagenesis experiments (Supplementary Fig. 1A, B). Pilot studies were conducted with TILL8 seeds to balance a desired high mutation frequency with plant viability and fertility required to produce a sufficiently large number of M_1_ plants and M_2_ seed. Imbibition serves to activate seeds physiologically and reduce somatic effects of EMS treatment. To modulate EMS effects in the final mutagenesis treatments, the imbibition rate was calculated and revealed a fast increase of water uptake in the first 8 h until reaching a maximum of 50% at 24 h (Supplementary Fig. 1C). The effect of two durations of imbibition (0 and 2 hours) when seeds had reached either 20 or 32% water uptake were then analyzed in combination with two concentrations of EMS (150 mM, 200 mM) and two exposure periods (3 and 5 hours) on whole-plant, root and shoot dry weight; root and shoot length; and seedling development. Plant, shoot and root dry weight were significantly reduced with every combination of pre-soaking and mutagen exposure (*p* < 0.05) while root and shoot lengths were unaffected (Supplementary Fig. 2A-E). In examining seedling development compared to controls, a survival rate close to 50% (termed as lethal dose 50, LD_50_) was achieved combining 2 h of pre-soaking with 3 h of EMS at 200 mM concentration (Supplementary Fig. 2F). Using these preliminary experiments as a guide, the definitive, large-scale mutagenesis was carried out in two separate experiments using 2 h of pre-soaking combined with 3 h of EMS exposure on 1,000 seeds each at 150, 200 and 250 mM concentrations.

### Generation of a mutant population for TILLING by sequencing

The M1 mutant plants comprising the TILLCANN population were cultivated in two cycles to harvest seed from a total of 1,633 M2 families (Figs. 1 and 2A). Most of the M1 mutant plants used to produce M2 seed families were derived from either 200 mM or 250 mM EMS treatments (48 and 42% of the population, respectively) while only 10% were produced from seeds treated with 150 mM EMS (Table S1). Developmental abnormalities, embryo lethality, and alterations in leaf morphology were conspicuous mutant phenotypes detected in M1 plants and served as reliable visual indicators of successful mutagenesis (Figs. 2B - 2E). Another interesting mutant phenotype was an M1 plant that had reverted to partial monoecy, with a single branch producing late male flowers together with early female flowers (i.e. family 200mM_827).

**Figure 1.**
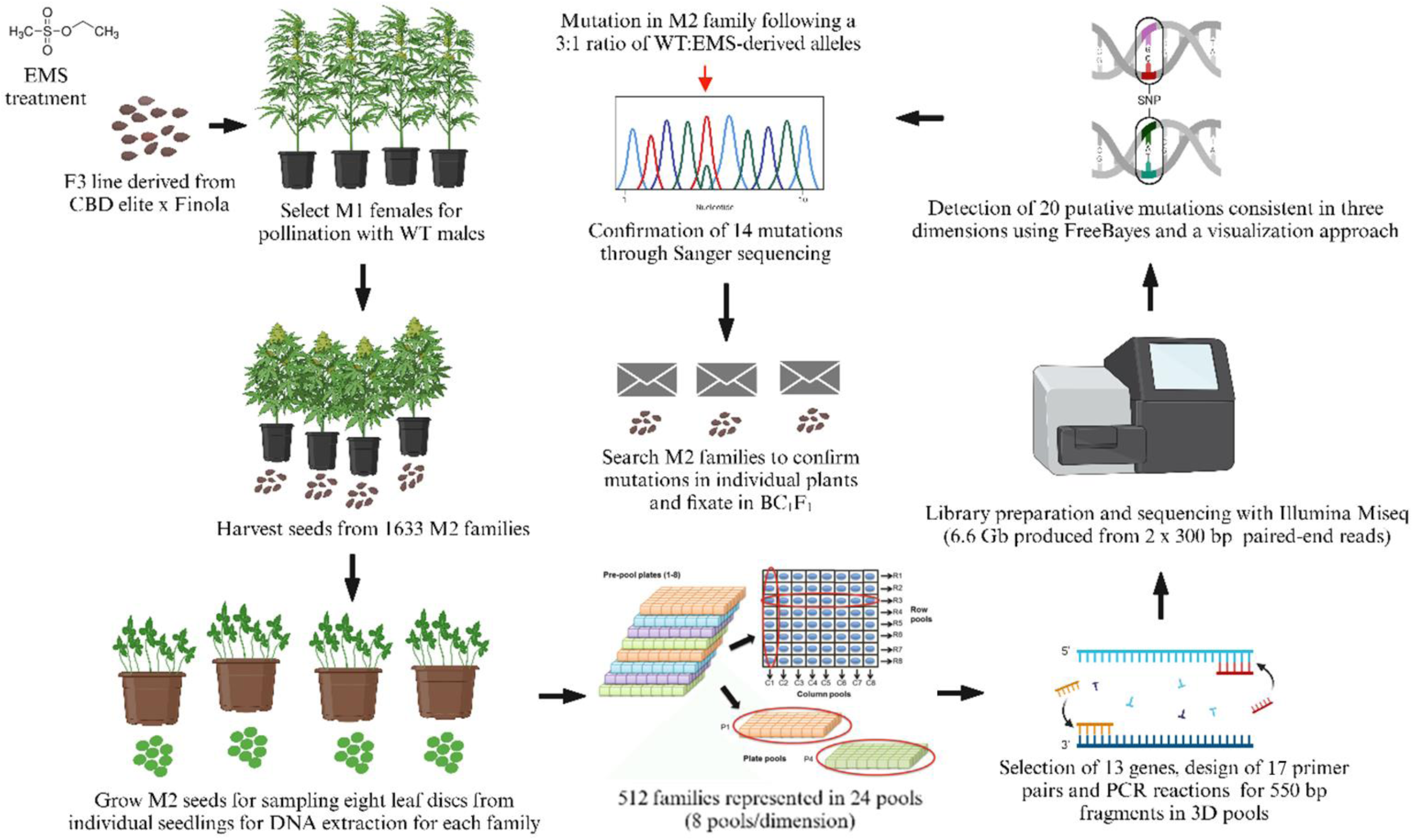
Workflow for the TILLING population construction and the detection of mutations in tridimensional pools. Created with BioRender.com. Tri-dimensional pooling diagram reproduced from Burkart-Waco et al. (2017) under Creative Commons license CC-BY-NC 2.5 (https://creativecommons.org/licenses/by-nc/2.5/).

**Figure 2.**
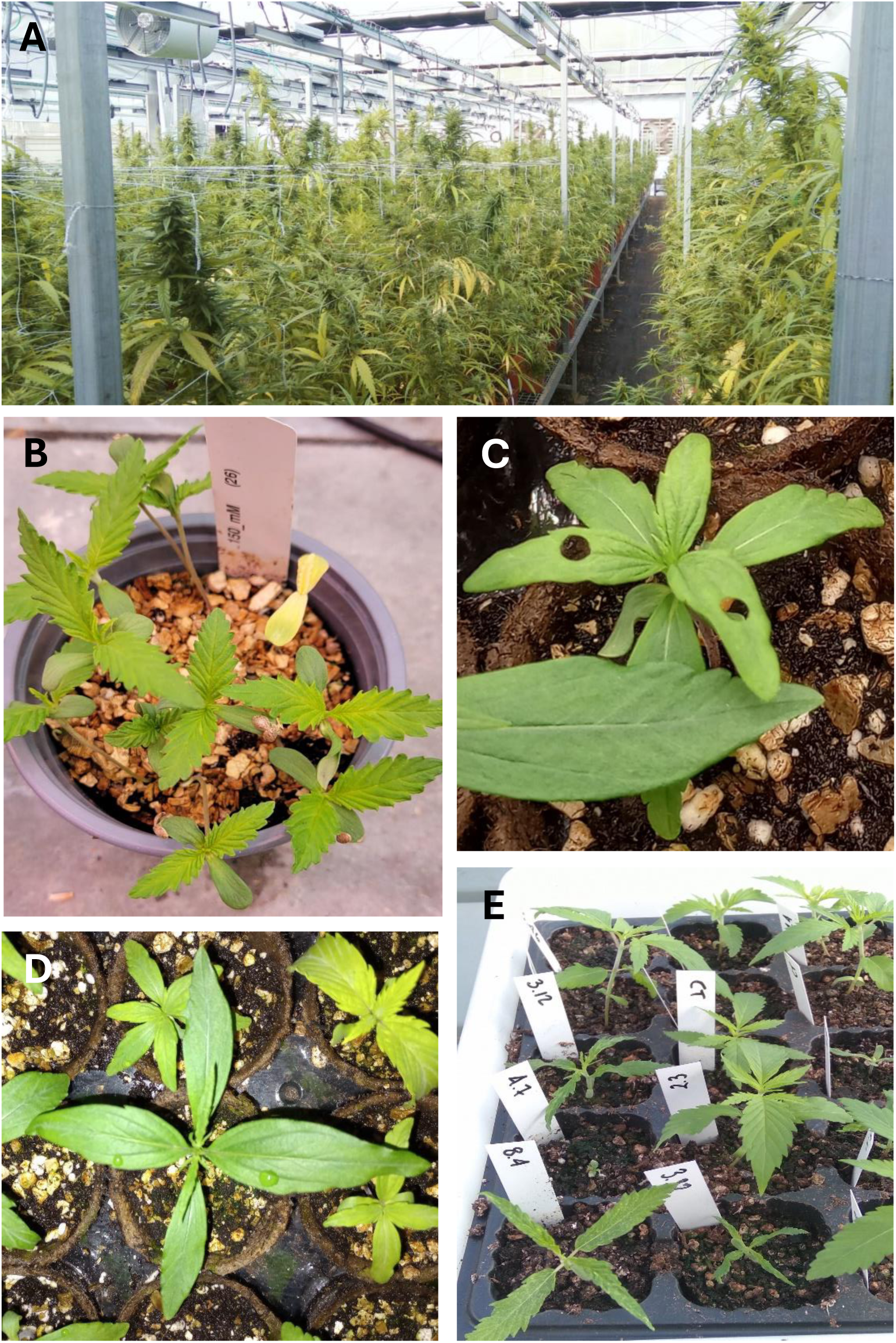
Phenotypic characteristics of the TILLCANN population cultivated in permitted greenhouses (A), mutant seedlings displaying chlorosis (B) alterations in cotyledon number and leaf phyllotaxis (C), altered leaf morphology (D), and developmental abnormalities (E).

### Mutations revealed by whole genome re-sequencing

The whole genome re-sequencing (WGRS) of six M2 plants were performed to assess the mutation frequency across the genome. Reads included in the mutant libraries showed an optimal mapping efficiency to the cs10 genome (90%), with accurate mate pairing (86%) (Table S2). Around 60% of the genome was covered in each library with a proper depth for SNP calling obtaining a mean depth of 40.1x ± 0.6 across the genome for the six libraries. After variant detection and filtering to retain canonical EMS mutations in positions indicated using WGRS data from F1 parental lines of the TILL8 population, we identified and annotated between 260 and 17,179 homozygous mutations per genome in the six mutants (Supplementary Figure S3; Table S3). Most of the putative mutations were intergenic (14,274) followed by variants located upstream from the start codon and the 5’-end of the gene in the promoter regions. The least frequent variants corresponded to those in exonic regions affecting protein function including missense, splice site and stop-gained mutations (804). Mutations tended to cluster at the beginning of chromosomes and in regions of higher GC content (Supplementary Figure 4). Considering the 23,614 canonical mutations, an ungapped genome size of 736 Mbp for cs10, and 60% genome representation (441 Mbp), a density of 1 mutation per 112 kb, equivalent to 8.9 mutations per Mbp and a total of 3,935 mutations per genome was calculated in the TILLCANN population

### TbyS and mutation detection

To assess our platform for the efficiency in obtaining mutations, we conducted TbyS on a subset of 512 M2 families using 17 PCR amplicons designed from 13 *C. sativa* genes (Fig. 1). Targets for mutation discovery included homologues for powdery mildew resistance (*CsMLO1*); flowering related genes apetala 2 (*CsAP2),* flowering time (*CsFT)*, agamous-like 6 (Cs*AGL6)*; sex determination giberrellin insensitive (Cs*GAI*); cannabinoid biosynthesis and cannabidiolic acid synthase (*CsCBDAS)*, 3-oxoacyl-[acyl-carrier-protein] reductase 4 (Cs*BKR)*, and two copies of olivetol synthase (Cs*OLS-1* and *CsOLS1-2*) and prenyltransferase 4 (*CsPT4)*; and for trichome formation including two copies of *CsTCP4* as well as *CsMYB106* (Table 1). Most gene targets co-localize with QTL or were selected based on orthology and functional validation in other crop species. Amplicons encompassed conserved genomic regions within exons with sizes ranging from 452 to 704 bp (Table S4).

**Table 1.** Genes selected to detect mutations with putative effect on agronomic traits and cannabinoid biosynthesis in a 3D-pooled cannabis TILLING population using an Illumina Miseq platform. The position in the chromosome (Chr) is according to cs10 reference genome.

| Gene | Loci cs10 | Functional annotation | Chr | Trait |
| --- | --- | --- | --- | --- |
| *CsMLO1* | LOC115705883 | Mildew resistance locus 1 | 1 | Powdery mildew response |
| *CsAP2* | LOC115708151 | Floral homeotic protein<br>Apetala 2 | 1 | Flowering time |
| *CsFT* | LOC115700781 | Florigen locus, PEBP family | 8 |  |
| *CsAGL6* | LOC115704294 | Agamous-like MADS-box protein | 1 | Flowering time, sex determination |
| *CsGAI* | LOC115711473 | GRAS-type transcription factor (TF) | 3 | Sex determination |
| *CsBKR* | LOC115714062 | 3-oxoacyl-[acyl-carrier-protein] reductase 4 | 4 | Cannabinoid biosynthesis |
| *CsCBDAS* | LOC115697762 | Cannabidiolic acid synthase | 7 |  |
| *CsOLS1-1* | LOC115699293 | Olivetol synthase | 8 |  |
| *CsOLS1-2* | LOC115700696 |  |  |  |
| *CsPT4* | LOC115713185 | Aromatic prenyltransferase | X |  |
| *CsTCP4-1* | LOC115697611 | Class II TEOSINTE BRANCHED 1/CYCLOIDEA/PCF TF | 7 | Trichome formation |
| *CsTCP4-2* | LOC115600698 |  | 8 |  |
| *CsMYB106* | LOC115701410 | MIXTA-like MYB gene NOECK (NOK) TF | 8 |  |

To perform amplicon sequencing, first, DNA from each of the 512 M2 families was pooled tridemensionally (i.e. eight pools per column (C), row (R) and plate dimensions (D)) to generate 24 DNA pools representing 64 families per pool). Sequencing of the 17 PCR amplicons in each of the pools produced 888,094 reads/pool on average (Table S2). Mapping efficiency was >94% and 90.8% of reads were properly paired during the mapping. The percentage of reads with a mate mapped to another chromosome/amplicon was low (0.7%). Reads from C and R pools mapped equally efficiently, while more variability was present in D pools. Mean sequencing coverage of amplicons was 11,584 (11.3x per M2 family) and was highly variable, ranging from 1.1 – 50,000 reads/amplicon (Table S5). Sequencing coverage of amplicons was also highly variable between pools and most amplicons mapped preferentially either to the cs10 or Finola genomes, evidenced by low mean coverage (< 2,000 reads) when mapping to the non-preferential genome. For example, reads from amplicons for AP2_F2R1 mapped efficiently to the Finola genome, and poorly to cs10, with a mean coverage across pools of 12,608 reads in the former compared to 174 reads in the latter (Fig. 3). This was also true for OLS1-1_F2R2 and TCP4-2_F2R2, while the opposite was the case for MYB106_F2R2, OLS1-1_F1R1 and GAI_F2R2, which mapped more efficiently to cs10. Additionally, some amplicons failed completely to map in one genome vs the other, with BKR_F1R1 and FT mapped exclusively to cs10, while Mlo1_F3R2 mapped exclusively to Finola. Despite this variability, sequencing was of sufficient depth in all cases for amplicons to be analyzed for variants in the preferential genome where the reads were mapping. In contrast, reads for GAI_F1R1 failed to map adequately in either genome so this amplicon was excluded from subsequent variant calling. As each pool in the C, R, and D dimensions contains 64 families x 8 plants per individual, the 1024 alleles present were sequenced to a depth from 2.2x to 48.9x.

**Figure 3.**
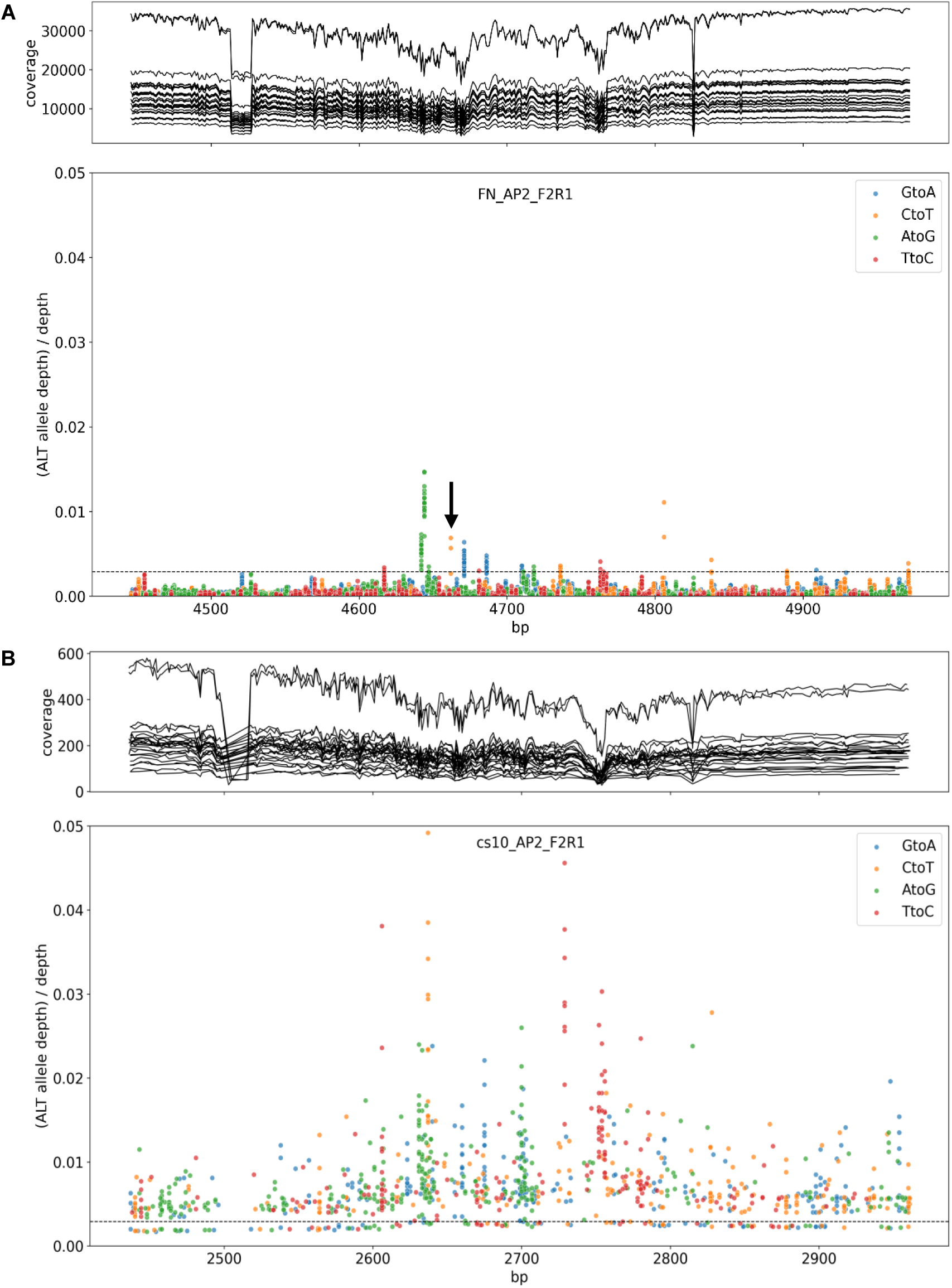
Detection of putative EMS-derived mutations based on per-base sequencing coverage in each of 24 3D-pooled libraries in which 512 M2 families were represented (solid lines, top panel) and per-base frequency of the alternative alleles at gene positions (bottom panel) for amplicon AP2_F2R1 mapped to the Finola (A) and cs10 (B) genomes. Solid arrow represents EMS canonical transitions for a mutation at position 4,662 bp in three libraries which represent the three outlier pools with putative mutations exceeding or close to 0.0029 (dashed line, bottom panel).

Following the SNP calling procedure using FreeBayes (Garrison and Marth, 2012) and filtering to retain low frequency variants, the detection pipeline identified 29, 31 and 31 variants in C, R, and D pools, respectively, at the threshold limit of detection (mutant allele frequency = 0.0025) (Supplementary Fig. 5). Five G:A mutations were consistent in all dimensions and were characterized as very high probability mutants. Six variants (five C:T and one non-canonical A:C) were detected in two dimensions and classified as likely mutants. Alternative allele frequencies were called at the positions of these variants to infer the pools from the additional dimension containing the SNPs and subsequently, the M2 families carrying the mutations. Nine additional mutations were identified through a visualization approach based on the scoring of the alternative allele frequencies exceeding or close to the limit of detection at each position across the amplicons (Fig. 3). Putative mutations were pinpointed when three different pools distributed in the three dimensions showed alternative allele frequencies in the vicinity of the threshold limit of detection. Using these combined approaches, 20 putative mutations were detected in 22 M2 families. Two families accumulated two mutations in different genes (i.e. 200mM_114 and 250 Mm_240) and two mutations were predicted ambiguously in D pools for 250mM_75 and 200mM_228 in amplicon MYB106_F2R2 and in R pools for 250mM_249 and 150mM_7 in amplicon BKR_F1R1 (Table S6).

### Confirmation by Sanger sequencing and PACE marker development

To confirm putative mutations in the identified M2 families, Sanger sequencing reads were produced for the amplicons from the DNA samples used to perform the 3D pooling. Fourteen canonical EMS-derived mutations for eight genes in 12 different families were validated through Sanger sequencing (Table 2). Three mutations were not observed at predicted positions and three other mutations could not be confirmed as trace files contained signal noise in the sequence flanking the variant due to indel polymorphisms (Table S6). Overall, three mutations were confirmed in *CsAP2*, three mutations in *CsOLS1-1* and two in *CsOLS1-2* two mutations in both *CsMYB106* and *CsTCP4*, and one mutation each in *CsBKR*, *CsPt4*, and *CsMLO1*. The two mutations in *CsTCP4* were in the same position in the gene model from cs10 and Finola and were found in two independent families. Eight of the confirmed mutations were observed in families originating from the 200_mM EMS treatment, the concentration with highest frequency for the families included in the 3D-pooling design. Four mutations originated from the 250_mM treatment, while two came from the 150_mM treatment. By sequencing 8,739 bp contained in 17 amplicons and multiplied by 512 families sequenced, we analysed 4,474 Mbp of total sequence yielding a mutation frequency of 1/320 kb based on 14 confirmed mutations out of 17 that could be accurately analyzed by Sanger sequencing.

**Table 2.** List of Sanger-confirmed EMS-derived mutations detected in pools and families with the predicted effect on protein function in cs10 and Finola.

| Amplicon and primer combination | Chr | Mutation and gene position<sup>a</sup> | C pools |  | D pools |  | R pools |  | M2 Family | Predicted effect<sup>e</sup> | Prediction scores for SAASs<sup>f</sup> |
| --- | --- | --- | --- | --- | --- | --- | --- | --- | --- | --- | --- |
|  |  |  | Pools | Frequency | Pools | Frequency | Pools | Frequency |  |  |  |
| *MLO1_F1R1* | 1 | C<sub>6894</sub>T | C3 | 0.0022 | D7 | 0.0025 | R8 | 0.0056 | 250mM_274 | intron mutation |  |
| *AP2_F2R1* | 5 | C<sub>4806</sub>T | C2 | 0.0070 | D2 | 0.0022 | R6 | 0.0111 | 200mM_336 | synonymous mutation A |  |
| *AP2_F2R1* | 5 | C<sub>4662</sub>T | C8 | 0.0027 | D8 | 0.0069 | R7 | 0.0057 | 150mM_9 | synonymous mutation I |  |
| *AP2_F2R1* | 5 | C<sub>4838</sub>T | C1 | 0.0028 | D8 | 0.0043 | R3 | 0.0029 | 200mM_120 | A139V neutral | 0.023 |
| *TCP4-1_F2R2* | 7 | C<sub>3838</sub>T | C1 | 0.0074 | D7 | 0.0055 | R7 | 0.0090 | 250mM_240 | P163S neutral | 0.008 |
| *TCP4-1_F2R2* | 7 | C<sub>3172</sub>T | C2 | 0.0031 | D4 | 0.0069 | R5 | 0.0016 | 200mM_99 | P163S neutral | 0.032 |
| *BKR_F2R2* | 7 | G<sub>4044</sub>A | C1 | 0.0353 | D7 | 0.0094 | R7 | 0.0221 | 250mM_240 | synonymous mutation L |  |
| *OLS1-1_F1R1* | 8 | G<sub>2761</sub>A | C8 | 0.0153 | D7 | 0.0203 | R3 | 0.0108 | 250mM_201 | A167T functional | 0.999 |
| *MYB106_F2R2* | 8 | C<sub>3779</sub>T | C7 | 0.0090 | D1 | 0.0016 | R2 | 0.0102 | 150mM_29 | synonymous mutation T |  |
| *MYB106_F2R2* | 8 | C<sub>3745</sub>T | C7 | 0.0035 | D2 | 0.0009 | R1 | 0.0039 | 200mM_228 | A258V neutral | 0.007 |
| *OLS1-2_F1R1* | 9 | G<sub>2827</sub>A | C6 | 0.0140 | D1 | 0.0129 | R5 | 0.0070 | 200mM_119 | synonymous mutation L |  |
| *OLS1-2_F1R1* | 9 | G<sub>2975</sub>A | C1 | 0.0061 | D1 | 0.0022 | R3 | 0.0050 | 200mM_114 | P131L functional | 0.946 |
| *OLS1-1_F2R2* | 9 | C<sub>2649</sub>T | C1 | 0.0053 | D1 | 0.0032 | R3 | 0.0105 | 200mM_114 | P131L functional | 0.941 |
| *Pt4_F1R1* | X | G<sub>9952</sub>A | C6 | 0.0267 | D2 | 0.0205 | R6 | 0.0120 | 200mM_259 | intron mutation |  |

<sup>a</sup>Position of the mutation based on the gene sequence.
<sup>e</sup>The effect of single amino acid substitutions was predicted with the PPVED software (Guo et al., 2022) and the mutations were classified as neutral when an impact in the protein activity is not expected or functional when protein performance is projected to be altered.
<sup>f</sup>Prediction scores (*Ps*) for the single amino acid substitutions (SAASs) obtained with the PPVED software (Guo et al., 2022). Neutral: *Ps*<0.5; functional: *Ps*≥0.5.

The family 200mM_114 possesses mutations in the both copies of *CsOLS*, exhibiting a base change from C:T at position 2,649 in *CsOLS1-1* and G:A at position 2,975 in *CsOLS1-2* (Table 2). These mutations are predicted to produce P131L substitutions with a strong probability (prediction score (*Ps*) > 0.94) of producing a functional mutation, defined as likely possessing a new molecular function or a new pattern of gene expression. A second mutation in *CsOLS1-1* contained in family 250mM_201 with a change from G:A at position 2,761 resulting in a A167T protein substitution is predicted to have an even higher probability (*Ps* = 0.999) of producing a functional mutation. The remaining mutations in amplicon sequences were predicted to produce synonymous amino acids or single amino acid substitutions (SAASs) with neutral effects and low predicted impacts on protein function.

To begin assessing mutant genotypes for a phenotype that might be associated with variation in the target locus, we first reconfirmed the presence of the mutations observed in pooled DNA samples in individual M2 plants through Sanger sequencing of individuals in families 200mM_119 for *ols1-1*, 200mM_99 for *tcp4-1*, and 150mM-29 for *myb106* (Fig. 4A, C, and E). We backcrossed the heterozygous M2 mutants and then designed specific PCR Allele Competitive Extension (PACE) markers to genotype backcross 1 (BC_1_) M2 individuals segregating for target mutations. We found perfect correlation between heterozygous mutant or homozygous WT individuals identified by sequencing and their corresponding marker genotypes (Fig. 4B, D, and F). All markers fit the 1:1 segregation ratio expected for BC_1_M2 plants where genotypes are either heterozygous or homozygous for WT alleles.

**Figure 4.**
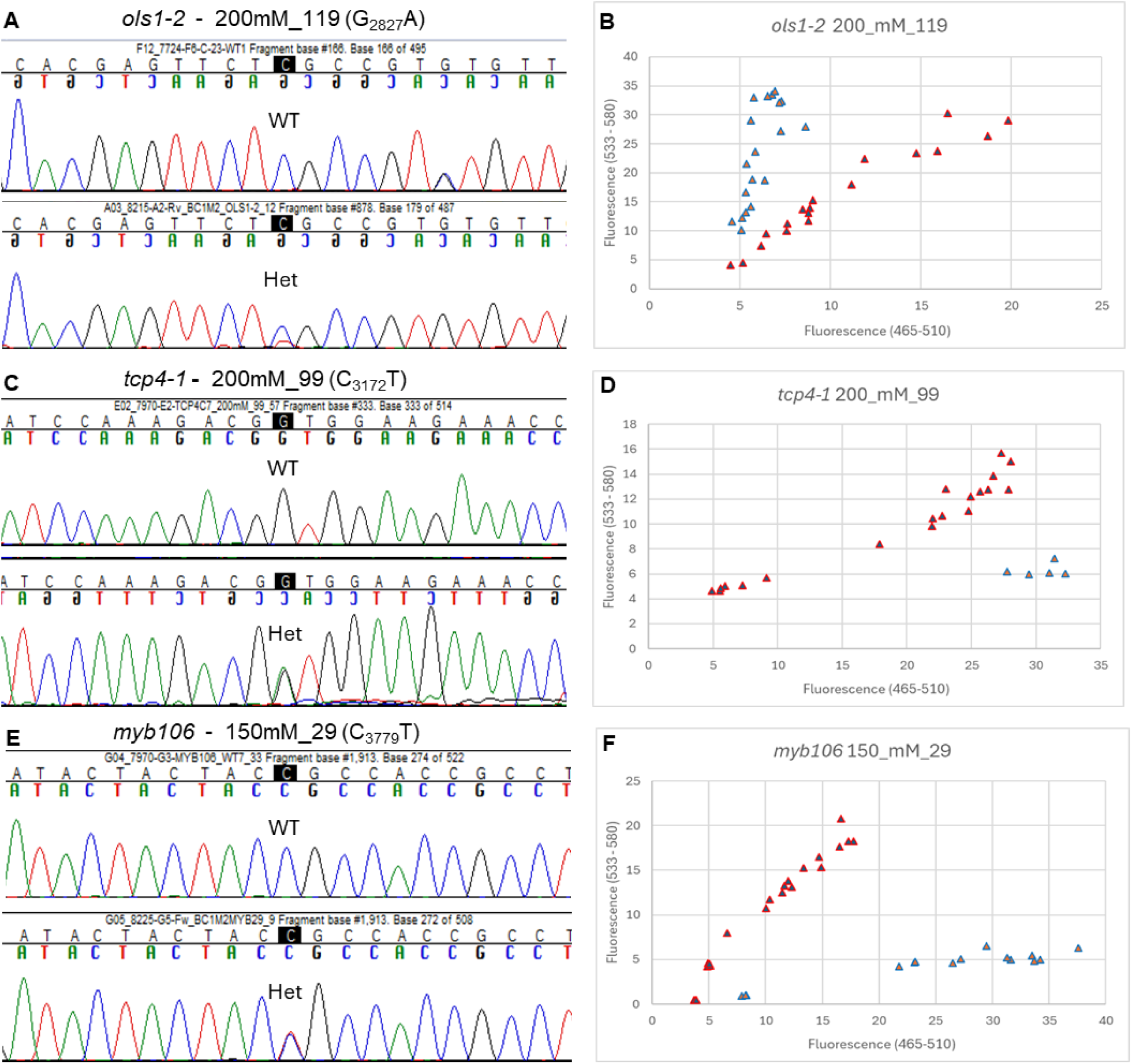
Verification of sequence variations (mutations) by Sanger dideoxy chain termination method in M2 mutant plants and conversion of sequence variants to PACE genotyping markers showing heterozygous individuals (red triangles) and homozygous (WT) individuals (blue triangles). The presence double peaks at shaded positions in a 1:1 ratio indicated the heterozygous state of the plant for mutations in families for genes *CsOLS1* at base position 2827 (A), and genotyping with PACE performed on 48 plants (B), mutation in *CsTCP4* at position 3172 (B) and genotyping results (D), mutation in *CsMYB106* at position 3779 (E), and genotyping result (F).

### Phenotyping of the *CsTCP4-1* mutation

We observed a striking phenotype in BC_1_ progeny associated with the mutation in *CsTCP4-1* contained in M2 family 200mM_99. When we grew out plants (n = 25), approximately half of the population showed pronounced phenotypic alterations in leaf number and morphology, having tri-foliate, glabrous leaves with reduced serration and venation (n=13) (Fig. 5A, C, D) compared to palmate, regularly serrated, rugous, leaves (n=12) typical of WT plants (Fig. 5B, D, E). These alterations showed complete linkage to the heterozygous mutant genotype, while individuals homozygous for non-mutant (reference) alleles were phenotypically WT (Fig. 4D). *CsTCP4-1* is composed of a single exon and the mutation results in a C:T base change at position 3,172 in the DNA sequence to produce a substitution of a serine for a proline at protein position 163 (Table 2). The PPVED mutation is predicted to produce a neutral substitution with a low effect (*Ps*=0.032). However, this mutation is located in a highly conserved sequence region spanning positions 158 – 168 (Supplementary Figure 6A) and is predicted by AlphaFold (Varadi et al., 2024; Jumper et al., 2021) to lie near the start of a helix spanning positions 167-178, albeit with a low per-residue model confidence score (pLDDT) of 50.75 (Supplementary Figure 6B). Introducing the mutation into the predicted folded protein structure substitutes an amino acid residue with an open ring and an additional exposed oxygen atom but fails to produce a conformational change in the protein (Supplementary Figure 6C, D).

**Figure 5.**
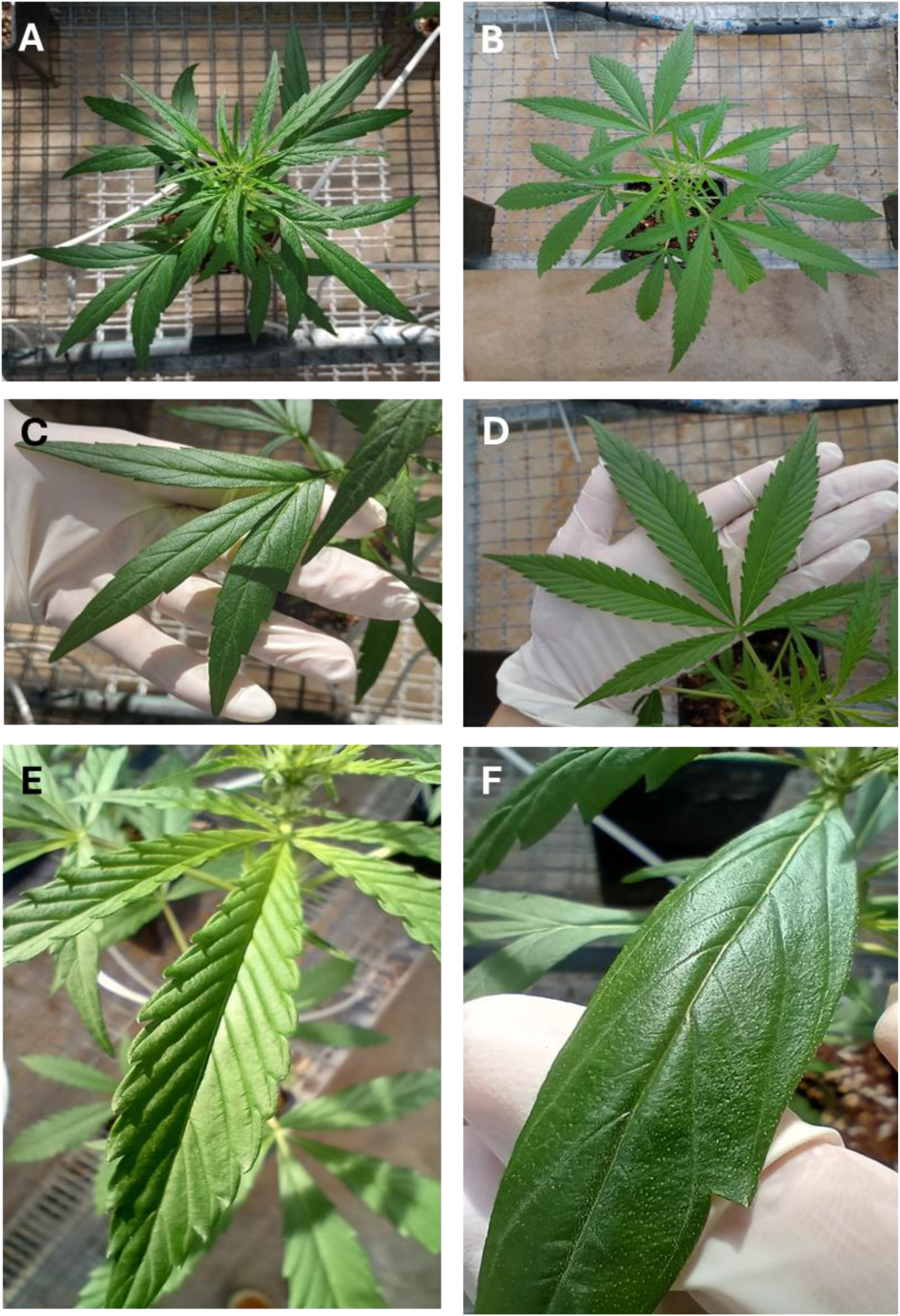
Phenotypic effects of the mutation of TCP4 on phyllotaxis for wild-type (A) and mutant (B), leaflet number for wild-type (C) and mutant (D), and leaf surface for wild-type (E) and mutant (F) plants.

## Discussion

### The TILLCANN platform as a high-quality resource for forward and reverse genetic screens

We developed and optimized a standard EMS mutagenesis protocol to create a TILLING population (TILLCANN) in *C. sativa* composed of 1,633 M2 families that is publicly available to the research community. As a first approach to determine the utility of this resource, we employed a high-throughput, TbyS strategy in a subset of 512 M2 families to identify and confirm fourteen canonical EMS-derived mutations for eight genes for important agronomic and biochemical traits (Table 1). Mutations were confirmed with 82% accuracy (14/17 producing quality Sanger sequencing results), which is within the range of values obtained in platforms for other species (Lakhssassi et al., 2021; Gupta et al., 2017; Jiang et al., 2022). Mutation frequency available in the TILLCANN platform is conservatively estimated at 1/320 kb, which is likely an underestimation due to three mutations that could not be unequivocally confirmed by Sanger sequencing. Therefore, a mutation frequency range of 1/263 - 1/320 kb is proposed to better represent the mutations that can be expected in exonic regions when screening for target genes of interest. As our platform represents the first publicly available TILLING resource in cannabis, we are unable to make direct comparisons. However, the calculated mutation frequency exceeds most mutagenized populations developed in other diploid species (reviewed in Szurman et al., 2023) and provides a high probability of recovering mutations in genes of interest. For example, based on a frequency of 1/263 or approximately four mutations per Mbp, a TbyS experiment using the full set of 1,633 M2 families in the TILLCANN platform would be expected to yield 8 mutants in a target gene, with a nearly 80% probability of recovering a deleterious missense mutation and an approximately 30% chance of finding a truncation (Heinkoff et al., 2004). Taken together, these results demonstrate the robustness and accuracy of mutant detection and verify the TILLCANN platform as a novel source for the creation of genetic diversity in cannabis. The platform is expected to be a valuable resource for the research community and will fill a role as in other important crops to provide new germplasm for basic studies of gene function and in breeding of improved commercial cultivars (Szurman-Zubrzycka et al., 2023). The population is versatile as the background from the F3 line that was mutagenized is derived from a cross between the industrial hemp line Finola and an elite Type III CBD-accumulating line that is low in THC. Therefore, it offers the possibility to screen mutations in specific genes for traits related to both industrial and medicinal applications. Improved genotypes with mutations introgressed from the TILLCANN platform can be used in traditional as well as organic agriculture systems according to European Union regulations (Bhattacharya et al., 2023). The market for cannabis-derived products with certifications related to social and environmental responsibility (fair trade, organic, etc.) is experiencing significant growth (Bennet 2021; Otañez, 2021) and cannabis sold in pharmacies is of mandatory organic cultivation (Baudean, 2021). Within this organic agriculture landscape, our platform can also play a role in the breeding of novel genotypes.

Data from WGRS showed that EMS treatments produced a mean of 3,935 mutations per line. However, just 5.3% of mutations were in coding regions, with 3.4% predicted to severely affect protein function. The ability to design amplicons in exons of target genes therefore makes TbyS an effective approach in cannabis to focus on these rare genic mutations. The mutation frequency of 1/112 kb using WGRS data was higher than the mutation frequency calculated by amplicon sequencing. Similar to results from Jiang et al (2022) the differences in the two estimates may be due to (1) false positives in mutation calling by whole-genome sequencing; (2) overstringency in variant detection or (3) underrepresentation of mutations in our targeted amplicons. Despite these differences, both the WGRS data and amplicon sequencing results supported a highly effective mutagenesis. Furthermore, mutation frequency calculated by WGRS data also provided insights into the mutational landscape of the genome by revealing that mutants are enriched with upstream variants present in promoter regions and that mutation frequency was influenced by adjacent sequence context (Supplementary Fig. 4) which will be useful in future studies with cannabis.

The frequency of mutations other than canonical transitions varies widely across species (Gupta et al., 2024) but is typically >90% in TILLING populations. In the TILLCANN population, all of the discovered mutations confirmed by Sanger sequencing corresponded to G:C to A:T base transitions typically induced by EMS-mutagenesis. Considering that the generation of TILLING populations in out-crossing species requires extensive precautions compared to requirements in autogamous species (Fanelli et al., 2021) a high degree of confirmed canonical mutations likely indicates a low level of genetic contamination in our population.

A proper depth of pooling is a crucial factor required for precise variant calling for a TbyS project where mutant allele frequency should exceed 0.0039 at a 1:256 pooling depth to rise above the sequencing noise in amplicon pools (Gupta et al., 2017). To increase sensitivity of mutant detection, we further decreased the threshold to >0.0025 to account for scenarios of obtaining only three mutant alleles per pool, or in cases where mutant containing reads may have been discarded due to poor quality. By combining two analysis pipelines (FreeBayes and visual inspection by plotting alternative allele frequencies across amplicons), we were able to overcome sequencing coverage heterogeneity across pools and amplicons to identify many mutations using this lower frequency threshold. The lower limit for mutant detection and confirmation was 0.0009 for MYB106_F2R2 while minimum allele coverage for a confirmed mutant was 2.2x for BKR_F2R2. These results suggest a possible lower limit of detection can be used at 1:256 pooling depths to capture mutations and that future TbyS studies in cannabis can use larger mutagenized populations to increase efficiency by potentially increase pooling depths while maintaining similar sequencing outputs.

Genetic studies in cannabis support a division between hemp and drug-type Cannabis (Schwabe et al., 2021; Ren et al., 2021). As a cross between these two lineages, TILL8 has an admixed genome. This can complicate read mapping efficiency and diminish variant detection as some portions of sequencing reads would not be expected to map correctly using only cs10 as the reference genome (considered drug-type, sharing 89% of its genome with marijuana varieties) (Grassa et al. 2021). In fact, we observed a high % of reads that mapped preferentially to the cs10 or Finola genomes and that some reads could only be unambiguously assigned to one of the genome variants for the corresponding amplicons. Mapping reads to two genomes therefore increased the probability of mutant detection through the augmentation of read mapping efficiency by decreasing mapping ambiguity (Tables S2, S5). One advantage is that with this approach we were able to discover redundant mutations located separately in the Finola and cs10 genomes in two independent families for *CsTCP4* that by only considering one reference genome would have been omitted. This was offset by the fact that two amplicons from Finola and cs10 were identical and reads mapped ambiguously in their respective regions. In this case, variants could not be scored in regions with reads with these ambiguous alignments, and variant detecting power was sacrificed.

### Phenotypic validation and predicted effects of induced mutations in candidate genes for trichome formation and cannabinoid biosynthesis

The TCP gene family is involved in the multilayer control of leaf development (Koyama, 2017) and has roles in the suppression of trichome initiation and trichome cell branching (Vadde et al., 2019; Vadde et al., 2018). The mutation in *CsTCP4* contained in the M2 family 200mM_99 was predicted by PPVED software to produce a neutral amino acid change and a low impact on protein function. However, functional consequences of conservative mutations can be difficult to predict (Fowler and Fields, 2014) which was evidenced in our study. We found perfect co-segregation between the trifoliate/glabrous/non-serrated leaf phenotype and the *tcp4-1* mutant in heterozygosis (Fig. 5B, D, F) supporting a strong causal relationship for the involvement of *CsTCP4-1* on wide range of phenotypic effects on cannabis leaf number and morphology. This result provides a validation of the TILLCANN platform as a resource for producing novel sources of genetic and phenotypic variation while reinforcing the idea that even non-target mutations or those predicted to have little to no impact can unlock interesting phenotypic or biochemical diversity when assessed and are worth exploring to diversify traits in cannabis. Although it didn’t produce a change in protein folding as predicted by AlphaFold, the *tcp4-1* mutation is located in a conserved sequence region with predicted low confidence at the start of a helix, thus may still produce a conformational change in the protein. To have a deeper understanding on *CsTCP4* function and targets, a comparative DNA affinity purification sequencing analysis (DAP-seq) can discover TF-binding sites to reveal gene targets regulated differently in the WT and mutant genotypes in a fast and inexpensive manner (Bartlett et al., 2017). This information can be coupled with a transcriptomic analysis to observe genes with differential expression in the WT and *tcp4-1* homozygous mutants.

Besides *TCP4-1*, six other identified mutations produced SAASs in our targeted candidate genes, with three predicted to produce neutral effects (Table 2). Three others were functional mutations with high *Ps* identified in *CsOLS1* and two of them were those detected in the family 200mM_114 in the same position of the two paralogs. *CsOLS1* belongs to the type III polyketide synthases (PKS) family involved in the cannabinoid biosynthetic pathway producing olivetolic acid (OA) as a precursor to CBDA and tetrahydrocannabinolic acid (THCA) (Taura et al., 2009). SAASs in *CsOLS1* are in highly conserved residues identified through the comparison with protein sequences from 22 type III PKS family genes, including sequences from three bacteria and one fungi (Abe and Morita, 2010). The SAAS P131L is in the vicinity of the residues 132 and 133 lining the enzyme active site, while A167T is proximate to C164 which is an amino acid from the catalytic triad (Abe and Morita, 2010). Targeted mutagenesis of other active site residues in OLS produce alterations in OA concentrations (Kearsey et al. 2020). Therefore, functional mutations in *CsOLS1* are promising for their potential to both augment cannabinoid content for medicinal applications, or diminish it for industrial hemp applications where exceeding limits of THC production are a concern. Current work is focused on backcrossing to reduce background mutation loads and the confirmation of mutant phenotypes in segregating populations *ols1, pt4,* and *myb106*. The backcrossing procedure can be accelerated using “speed breeding” (Schilling et al., 2023) combined with genomic background selection (Karunarathna et al., 2021) to deliver pre-breeding germplasm in a more efficient way.

Large quantities of seed for most M2 families contained in the TILLCANN platform are available for study. This opens the possibility of performing relatively easier forward genetic screens requiring larger populations of seeds for tolerance to salt, water stress and the stress-signalling phytohormone abscisic acid by germinating seeds on media containing the stressors (Manzanares et al., 2016). These objectives fit squarely into the value proposition of cannabis produced for industrial hemp and medicinal/recreational purposes as a sustainable and environmentally friendly crop suitable for the circular economy by improving abiotic stress tolerance, phytoremediation capacity, or by reducing agronomic inputs and energy requirements required for cultivation.

## Material and Methods

### Optimization of mutagenesis conditions

The EMS mutagenesis was defined in F3 seed derived from the cross of an elite CBD-producing female and the hemp genotype Finola. The germination rate, seedling development and number of seeds from three F3 lines was assessed to select one line for massive mutagenesis. To define a duration for imbibition for the seeds, the imbibition rate was observed for 24 h and calculated as the percentage of water intake of seeds hourly as determined by an increase in seed weight after every hour of imbibition according to Unan et al. (2022).

After the imbibition rate study, a preliminary experiment was conducted to define the mutagenesis conditions. Two pre-imbibition times (0 and 2 h) and two mutagen exposures (3 and 5 h) were combined with two EMS concentrations (150 and 200 mM). EMS dilutions were prepared in 2% dimethyl sulfoxide (DMSO) and 0.3 mL/seed of solution was added to 100 seeds per treatment. Control treatments with distilled water and 2% DMSO were used. Seeds were surface sterilized with Captan 80% (2.5 g/L) fungicide for approximately one minute and then rinsed with tap water. Seeds were then imbibed in bottles with EMS solutions and placed on a rotary shaker at room temperature. The mutagenesis was inactivated by treating the seeds with sodium thiosulfate 100 mM in two rounds of 20 min in the rotary shaker. The seeds were then rinsed gently with tap water and distributed in Petri dishes (20 seeds/plate) with wet filter paper to assess seedling survival. Four days after mutagenesis, seedlings were transferred to trays with a substrate of peat moss, vermiculite and perlite in a 3:1:1 ratio in a growth chamber at 22 °C and 18 h of light. A completely randomized design was established to sample five plants per treatment after 21 days to score height and dry weight of roots and shoots. A one-way analysis of variance was used to determine if the effect on growth-related traits from at least one EMS treatment was different from the others. The EMS treatments were compared against the DMSO control with pairwise comparisons considered significant after obtaining t-tests with Bonferroni adjusted *p-values* < 0.05. These analyses were carried out using RStudio v 2024.04.0 (RStudio Team, 2024).

For development of the TILLCANN population, 2 h of imbibition combined with 3 h of EMS exposure on 1,000 seeds each at 150, 200 and 250 mM was carried out. EMS was inactivated as described above. Following inactivation seeds were germinated in wetted rolled paper towels stored vertically in sealed plastic containers in a chamber at 24 °C with 18 h of light. The mutagenesis was repeated twice to coincide with space limitations and desirable cultivation conditions in the greenhouse to produce M1 plants and M2 seed.

### Construction of the TILLING population

At four days post-mutagenesis, seedlings were transferred to the greenhouse at CRAG and planted in 48-well trays with biodegradable pots filled with the substrate mix described previously. To maximize the probability of finding mutations, we prioritized transplanting seedlings from the 200 and 250 mM treatments. Males were visually discarded when floral primordia started to be conspicuous. Then, mutagenized females were transported to the IRTA research station in Caldes de Montbui (Spain; 41°37′54′′ N - 2°10′0.73″ W) and transplanted in 13 L plastic pots at 30 days post-mutagenesis. The pots were distributed in five elevated tables placed in a greenhouse under natural light conditions for the first experiment (harvested October 2022) and with supplementary light during the second experiment (harvested May 2023).

An M2 open-pollinated population was generated through the crossing of M1 females with wild-type males. Pollination was performed continuously for six weeks by shaking ten males across the plot with mutagenized flowering females. After ripening, the harvest of M2 families was performed by collecting the seeds from branches in individual trays using manual threshing. Seeds were deposited in brown kraft envelopes for drying at 23 °C and 45% of relative humidity for two weeks then stored at 4 °C under silica gel for long-term preservation.

### Mutation frequency assessed through WGRS

The mutation frequency of the TILLING population was assessed at the whole-genome level through the WGRS analysis of six M2 plants derived from different families subjected to two EMS concentrations (i.e. 200mM_103, 200mM_219, 200mM_227, 250mM_57, 250mM_231, 250mM_247). DNA was extracted from 100 mg of young leaves grounded with liquid nitrogen using the DNeasy Plant Pro Kit (Qiagen, Hilden, Germany). After isolation, DNA was quantified using a Nanodrop 2000c spectrophotometer (Thermo Fisher Scientific, Waltham, MA, USA) and the integrity was visually assessed in 1.5% (w/v) agarose gels stained with ethidium bromide. Tubes with 1.5 to 2.1 µg of DNA were sent to BGI Genomics (Shenzhen, China) for library preparation and sequencing. Libraries with insert size of less than 800 bp were sequenced with a MGISEQ-2000 platform (MGI Tech, Shenzhen, China) to produce 30 Gbp of 100 bp paired-end reads for each one. Thus, each mutant was sequenced with an expected 34x coverage.

Raw re-sequencing data were aligned to cs10 reference genome assembly (Grassa et al., 2021) (https://www.ncbi.nlm.nih.gov/datasets/genome/GCF_900626175.2/) using bwa v0.7.17-r1188 (Li and Durbin, 2009). We used Picard v2.22.3 (Broad Institute, 2019) to mark PCR duplicates and add read groups in each alignment file. Variant calling was performed using GATK (v4.1.7.0) (Van der Auwera & O’Connor, 2020) and Deepvariant (Poplin et a., 2018; Yun et al., 2021) with default parameters, selecting reads with minimum mapping quality higher than 10. After obtaining the raw variants from both callers, we selected those GATK variants that were also detected by Deepvariant. This common set of variants was filtered with hard filtering criteria, separately for SNPs and INDELs, as suggested by GATK best practices (https://gatk.broadinstitute.org/hc/en-us/articles/360035535932-Germline-short-variant-discovery-SNPs-Indels-) by using a minimum depth per-sample of 10, a minimum genotype quality of 20 and keeping sites with at least 90% of genotypes with data. To obtain the final list of putative EMS-derived mutations, a final set of three filters were applied to the variants. To detect a canonical EMS mutation in the WGS dataset we first selected positions with G<->A or C<->T transitions. Then we followed two strategies for getting the final list of putative canonical mutations. For the first strategy, we selected positions where both parents were heterozygous, five of the lines were heterozygous and one line was homozygous. This homozygous line would carryin theory the canonical mutation. For the latter strategy, we selected positions where both parents were homozygous, five of the lines were homozygous and the mutation-containing line was in heterozygosis. These variants were annotated with SnpEff (Cingolani et al., 2012).

### Detection of mutations by Illumina-based amplicon sequencing in a 3D-pooled TILLCANN population

The workflow for the detection of chemical mutations is shown in Fig. 1. A total of 512 M2 families composed of 319 families from the 200 mM treatment, 176 from the 250 mM treatment, and 17 from the 150 mM treatment were included for DNA extraction and TbyS. Seeds were grown in groups of 64 families distributed in 8 columns and eight rows of biodegradable pots (11 x 11 cm) at elevated tables in the CRAG greenhouse (20 °C mean temperature and 18 h light). Fifteen seeds per family were planted in the pots with the substrate described previously and covered with parafilm for 48 hours. Ten to 12 days after planting, one leaf disc of 6 mm diameter was collected from eight individual plants with a sharp hole punch. Then, eight discs from each family were homogenized with beads in a TissueLyser® disruptor (Qiagen, Hilden, Germany) to perform DNA extractions using the CTAB protocol (Doyle, 1991) with some modifications (Pereira et al., 2018). DNA quantification was performed with PicoGreen (Thermo Fisher Scientific, Waltham, MA, USA) using a Victor® Nivo™ (Perkin Elmer, Waltham, MA, USA) plate reader, followed by normalization to 20 ng/µl with a Mantis® liquid handler (Formulatrix, Bedford, MA, USA).

DNAs were pooled tri-dimensionally as described above to construct 24 DNA pools (C1-C8, R1-R8, D1-D8) and diluted to 5 ng/µl for PCR reactions. Pools were PCR-amplified using gene-specific primers. Owing to the hybrid nature of the genome of the TILLING population (i.e. a CBD-accumulating, cs10-like genotype and a fiber-use Finola genotype) conserved regions in both genomes were selected for primer design to produce consistent amplification of PCR products. Amplification conditions with KAPA HiFi HotStart Ready Mix (Roche, Basel, Switzerland) are described in the Supplementary Table S4 for each primer pair. Purified amplicons diluted in 52.5 µl of ultrapure water were quantified with PicoGreen and samples with concentrations greater than 0.3 ng/µl were used as template for Index PCR by merging for each sequencing pool 10 µl from each amplicon. The equal quantities of PCR products were combined in their respective pools, and combined amplicon pools were used for library preparation following the locus-specific primers protocol from the 16S metagenomic sequencing library preparation guide for the Miseq system (Illumina, 2013). Sequencing of libraries was performed on a MiSeq platform (Illumina, San Diego, CA, USA) that yielded 6.6 Gb of 2 x 300 bp paired-end reads.

Minimap2 (Li et al., 2018) was used to align the reads to a simplified reference file prepared with the sequence of the amplified genes retrieved from cs10 (Grassa et al., 2021) and Finola (Laverty et al., 2019) genomes. In each M2 family, a 3:1 ratio of WT:EMS-derived alleles are expected as 12 WT alleles were pooled with four mutated alleles in the tissue collection (i.e. four WT and four heterozygous plants with the mutated allele). Tri-dimensional pooling of DNA from the M2 families yields a pooling depth for mutation detection of 1:256 (four mutated alleles are expected in each pool with 512 families and 1024 alleles). Thus, the frequency change should rise above 0.0039 (4/1024) established as the detection limit to identify a mutation in an M2 family. We lowered this threshold to 0.0025 for the detection of variants using FreeBayes (Garrison & Marth, 2012) to account for scenarios of obtaining only three mutant alleles per pool, or in cases where mutant containing reads may have been discarded due to poor quality. The “pooled-continuous” option from the software was used to call variants simultaneously in the groups of libraries from each dimension. A mapping quality threshold of 20 was defined to consider reads for variant calling. After variant calling, a filter was applied to retain variants present in only one pool from each dimension. A constant of 2.8 was defined to multiply the alternative allele frequencies of the pools and distinguish putative chemical mutations from residual natural variability present as follows:

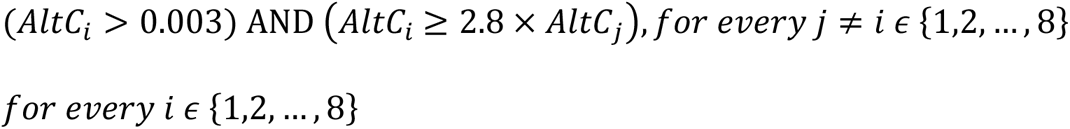

where *AltC_i_* corresponds to the alternative allele frequencies from the *i*th C pool. Variants consistent in three or two dimensions were identified. An additional pipeline was used to score further putative EMS-derived mutations and to identify the missing pool for the variants observed in two dimensions. Read depth at each base was plotted across the amplicons from the different pools to analyze the consistency of sequencing quality across the pools and identify additional mutants through visualization based on alternative allele frequencies that exceeded threshold detection limits at corresponding positions. Graphs were generated based on the alternative allele frequency of C:T, G:A, T:C, and A:G changes across the amplicon length for each library. The *mpileup* tool from Samtools (Li et al., 2009) was utilized to score read bases with phred-base quality 20 at each position, followed by the calculation of the frequency of the alternative alleles at each site. Subsequently, the graph revealed mutations at positions where three libraries from different dimensions showed an increased frequency of mutated nucleotides.

### Confirmation of mutations in M2 families and individuals through Sanger sequencing

The normalized DNA from the families diluted at 20 ng/µl was utilized in the PCR reactions for the confirmation of mutations through Sanger sequencing. Gene fragments were amplified using KAPA HiFi HotStart Ready Mix with the modification of some PCR conditions (Table S4). Sequencing was carried out at the Capillary Sequencing Facility from CRAG with an ABI 3730 DNA Analyzer (Applied Biosystems, Waltham, MA, USA) and fluorescent dye terminator detection. Examination of the sequence trace files was performed with Bioedit version 7.2.5 (Hall et al., 1999). These trace files were compared with the sequence of the unmutagenized genotype.

To reconfirm mutations in individual M2 plants, twenty-five seeds from unmutagenized line TILL8 and twenty-five seeds from each of the M2 families with functional mutations in *CsOLS1-1* and *CsOLS1-2* (200mM_114 and 250mM_201) as well as seeds of family M2 200mM_119 carrying a synonymous mutation in *CsOLS1-2*. We also germinated seeds of families 200mM_99, and 150 mM_29 for mutants *tcp4-1*, and *myb106*, respectively. Seeds were submerged in a prophylactic 80% thiram fungicide solution for approximately one minute and then rinsed with tap water. Seed were placed in petri dishes on saturated filter paper inside a translucent plastic box to maintain the humidity and incubated at 25 ℃ and 8 h of light. After three days, 16 developed seedlings for each family were transplanted and grown at 25 ℃ under 16/8 h diurnal cycle of light/darkness to maintain the plants in a vegetative state. DNA extractions, PCR, and Sanger sequencing were performed from young leaf tissue taken from each M2 plant as described previously to identify heterozygous female plants. After two weeks, lighting conditions were shifted to 12/12 h diurnal cycle to induce flowering. At flower initiation heterozygous female mutant plants from each family were backcrossed with TILL8 male plants and BC1M2 seeds harvested after six weeks.

### SNP marker development

To determine the zygosity of loci containing identified EMS-induced mutations, PCR Allele Competitive Extension (PACE®, 3CR Biosciences) markers were developed for each mutation in M2 plants as described previously (von Maydell, 2023) (Table S7). Primer Mix for each assay was prepared in final primer concentrations of 12 uM for A1, 12 uM for A2, and 30 uM for C1. DNA from M2 individuals with genotypes previously confirmed by Sanger sequencing were used to validate PACE® assays. Genotyping was performed using a LightCycler® 480 Instrument II (Roche), with detection type set to “Dual Color Hydrolysis Probe” and default PCR conditions. Results were analysed using LightCycler® 480 1.5.1.62 software. After validation, PACE® assays were conducted on BC1M2 DNA samples under the same conditions.

### Prediction of the effect of the EMS-derived mutations

The mutations were identified in either introns or exons according to the structural annotation of the genes in cs10 and Finola. The codons with the variants were considered to identify amino acid changes or synonymous mutations. The PPVED software is a machine learning tool designed for plants used to predict the effect of the SAAS on the protein function (Gou et al., 2022). A prediction score (*Ps*) from 0 to 1 is produced for the substitutions to classify them as neutral (i.e. *Ps*<0.5) when an impact in the protein activity is not expected or functional (i.e. *Ps*≥0.5) when changes in protein function are presumed. To assess the extent of sequence conservation across species, 50 full-length TCP4 homologues with the mutation were aligned using NCBI MSA Viewer 1.25.0. To visualize predicted protein structure of *CsTCP4*-1 and examine possible conformational changes induced by mutation, we used AlphaFold (https://alphafold.ebi.ac.uk/entry/A0A7J6HH57).

## Abbreviations

AGL6: agamous-like 6
AP2: apetala 2
PT: Aromatic prenyltransferase
BC: backcross
BKR: 3-oxoacyl-[acyl-carrier-protein] reductase 4
CBDAS: cannabidiolic acid synthase
CRISPR-CAS9: clustered regularly interspaced short palindromic repeats
cs10: CBDRx cannabis reference genome
CTAB: Cetyl trimethyl ammonium bromide
DMSO: dimethyl sulfoxide
EMS: Ethyl methanesulphonate
FT: flowering time
GAI: gibberellin insensitive
M: mutagenesis generation
Mbp: megabase pair
MLO: Mildew Resistance Locus O
mM: milimolar
MYB106: MYB106 transcription factor
NGS: Next generation sequencing
OLS: Olivetol synthase
PACE®: PCR Allele Competitive Extension
PKS: polyketide synthase
PPVED: Plant Protein Variation Detector
Ps: Prediction score
PT4: prenyltransferase 4
SAAS: single amino acid substitutions
SNP: Single Nucleotide Polymorphism
THCA: tetrahydrocannabinolic acid
TILLING: Targeting induced local lesions in genomes
TbyS: TILLING by sequencing
TCP4: Class II TEOSINTE BRANCHED 1/CYCLOIDEA/PCF
WGRS: whole genome re-sequencing
WT: wild type

## Supplementary data

**Supplementary Figure 1.** Germination (A) and seedling development (B) proportions for the three F3 lines assessed for large-scale mutagenesis experiments and (C) imbibition rate of seeds from TILL8 F3 line scored during 24 h.

**Supplementary Figure 2.** Effects of two EMS concentrations combining two imbibition times on plant, root, and shoot dry weights (A, B, C) and root and shoot lengths (D, E) and seedling development (F) scored under controls H2O and DMSO (2%). Asterisks represent significant differences of the EMS treatments with the DMSO control (p < 0.05) in the growth-related parameters affected by mutagenesis conditions according to analysis of variance.

**Supplementary Figure 3.** SnpEff annotation of canonical EMS-induced mutations detected by whole genome re-sequencing of six M2 plants.

**Supplementary Figure 4.** GC content (green line) in 5 Mbp sliding windows along ten cannabis chromosomes and counts of canonical mutations (bars) in 2 Mbp bins.

**Supplementary Figure. 5.** Venn diagram of the candidate mutations identified in each pooling dimension after the application of the alternative allele frequency filter.

**Table S1:** M2 families harvested for each EMS concentration during the two mutagenesis experiments (II-2022 and I-2023).

**Table S2**. Summary of mapping statistics for six whole genome re-sequencing (WGRS) and 24 amplicon-seq libraries (mean ± standard deviation).

**Table S3**. List of canonical EMS-derived mutations scored in six mutants analyzed through whole-genome re-sequencing.

**Table S4.** Functional insights of the genes selected for the detection of mutations, primer sequences and PCR conditions.

**Table S5**. Mean sequencing and allele coverage of amplicons sequenced in 24 DNA pools and mapped to the cs10 or Finola genomes

**Table S6.** List of EMS-derived mutations detected in pools and families with the predicted effect on protein function in cs10 and Finola (FN). Pools with frequency below 0.0025 are in red letters and highlighted in pink

**Table S7**. Genomic locations and sequences of primers used for PACE marker development

## Declarations

### Ethics approval and consent to participate

Research was conducted under permit from the Spanish Agency of Medicines and Medical Devices (AEMPS).

### Consent for publication

All authors approve the manuscript and consent to the publication of the work.

### Availability of data and materials

The main data supporting the results of this research are included within the article and the supplementary files provided. The files with raw reads from the 24 amplicon-seq and the six WGRS libraries were deposited to the European Nucleotide Archive (ENA) with study ID PRJEB81779.

### Author Contributions

DDD, AM, MP, CU, KA, AT and JA conceived and designed the study. RG, CZ, AM and JA acquired the funds for research and performed project administration. DDD performed mutagenesis experiments, constructed the mutant population, prepared the amplicon libraries and carried out bioinformatic analyses. NMA contributed to the bioinformatic analyses. VV designed PACE markers, made crosses, and validated the *tcp4-1* mutation. AM, CU, MP and JA supervised the research. KA supervised and contributed to the bioinformatic analyses. DDD and JA drafted the manuscript. All authors participated in editing the manuscript and approved the final version.

## Acknowledgments

We gratefully acknowledge the lab technical support provided by the staff of the Genomics and Biotechnology Laboratory at IRTA-CRAG and the Genomics Core Facility at CRAG.

