## Supplemental Figures for "TILLCANN: A TILLING platform in *Cannabis sativa* for mutation discovery and crop improvement"

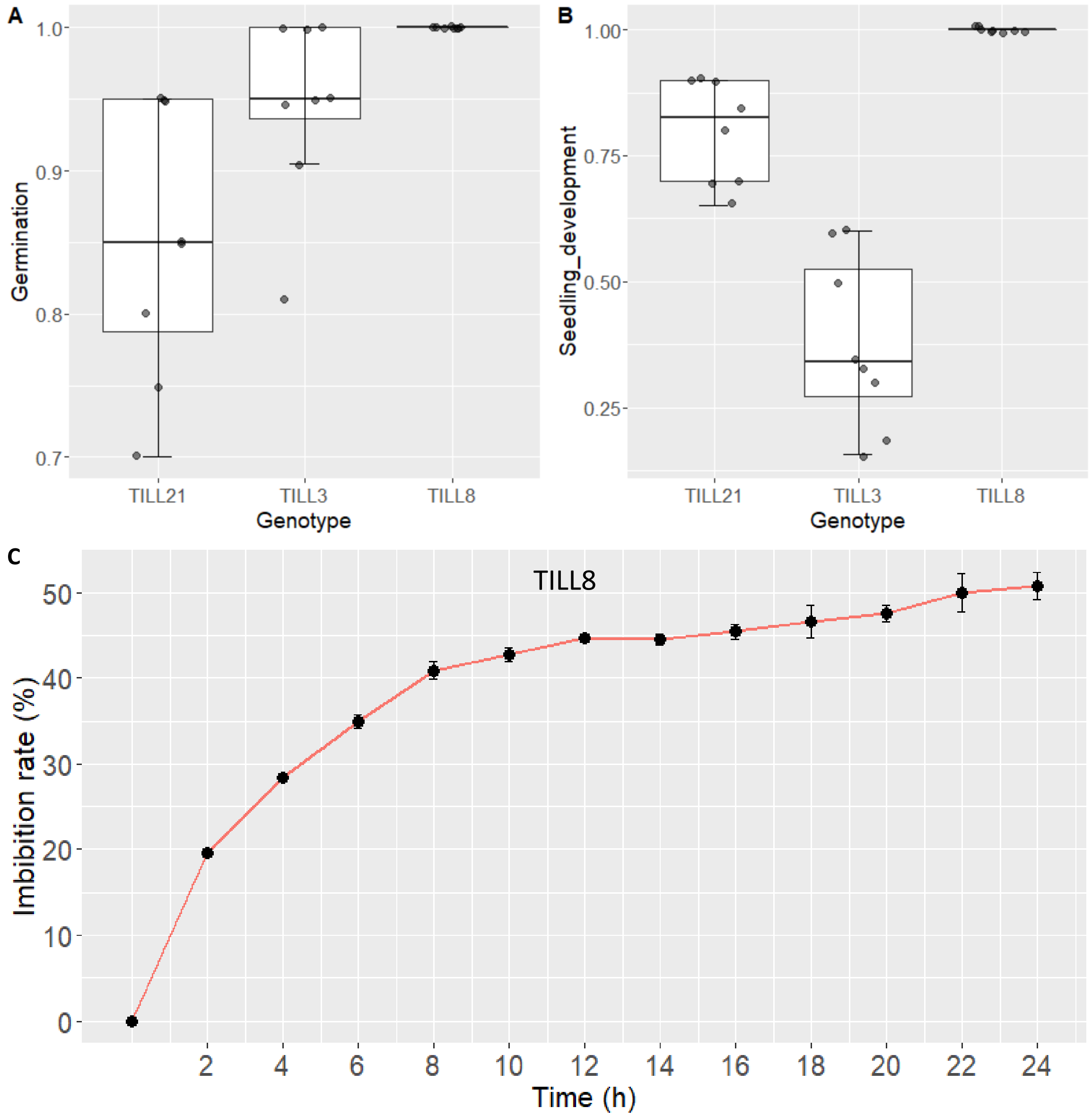


**Supplementary Figure 1.** Germination (A) and seedling development (B) proportions for the three F3 lines assessed for large-scale mutagenesis experiments, and imbibition rate for TILL8 (C).

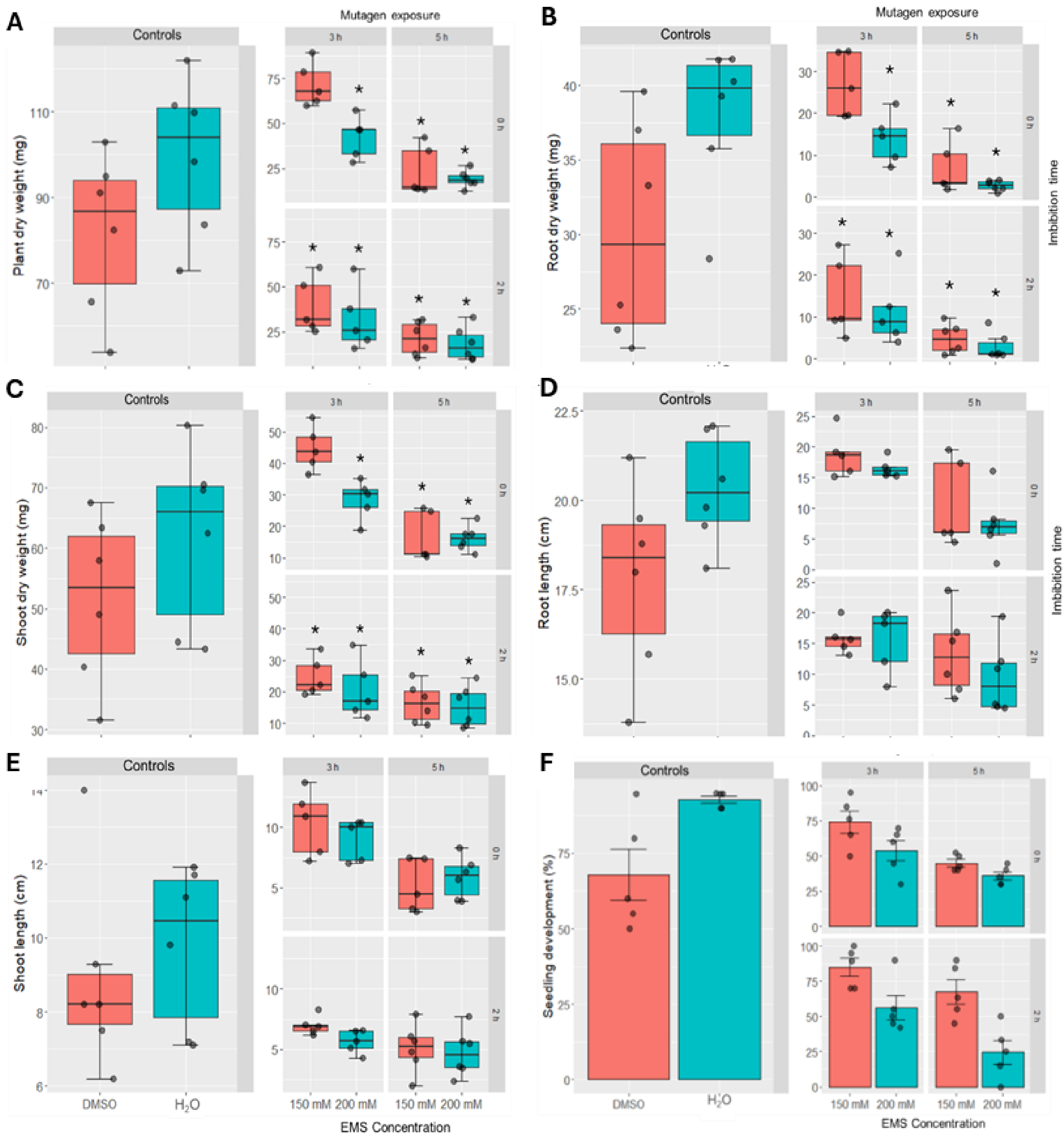


**Supplementary Figure 2.** Effects of two EMS concentrations combining two imbibition times on plant, root, and shoot dry weights (A, B, C) and root and shoot lengths (D, E) and seedling development (F) scored under controls $H_2O$ and DMSO (2%). Asterisks represent significant differences of the EMS treatments with the DMSO control ($p < 0.05$) in the growth-related parameters affected by mutagenesis conditions according to analysis of variance.

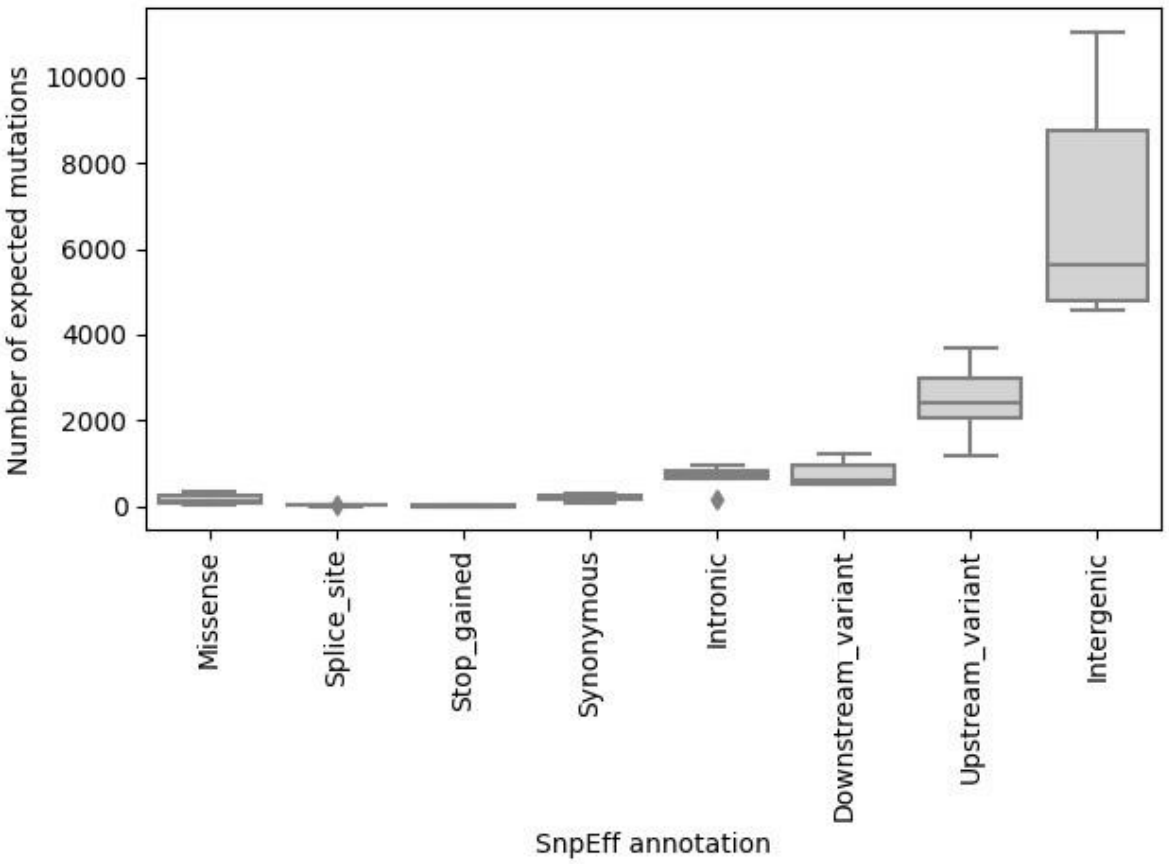


**Supplementary Figure 3**. SnpEff annotation (Cingolani et al., 2012) of canonical EMS-induced mutations detected by whole genome re-sequencing of six M2 plants.

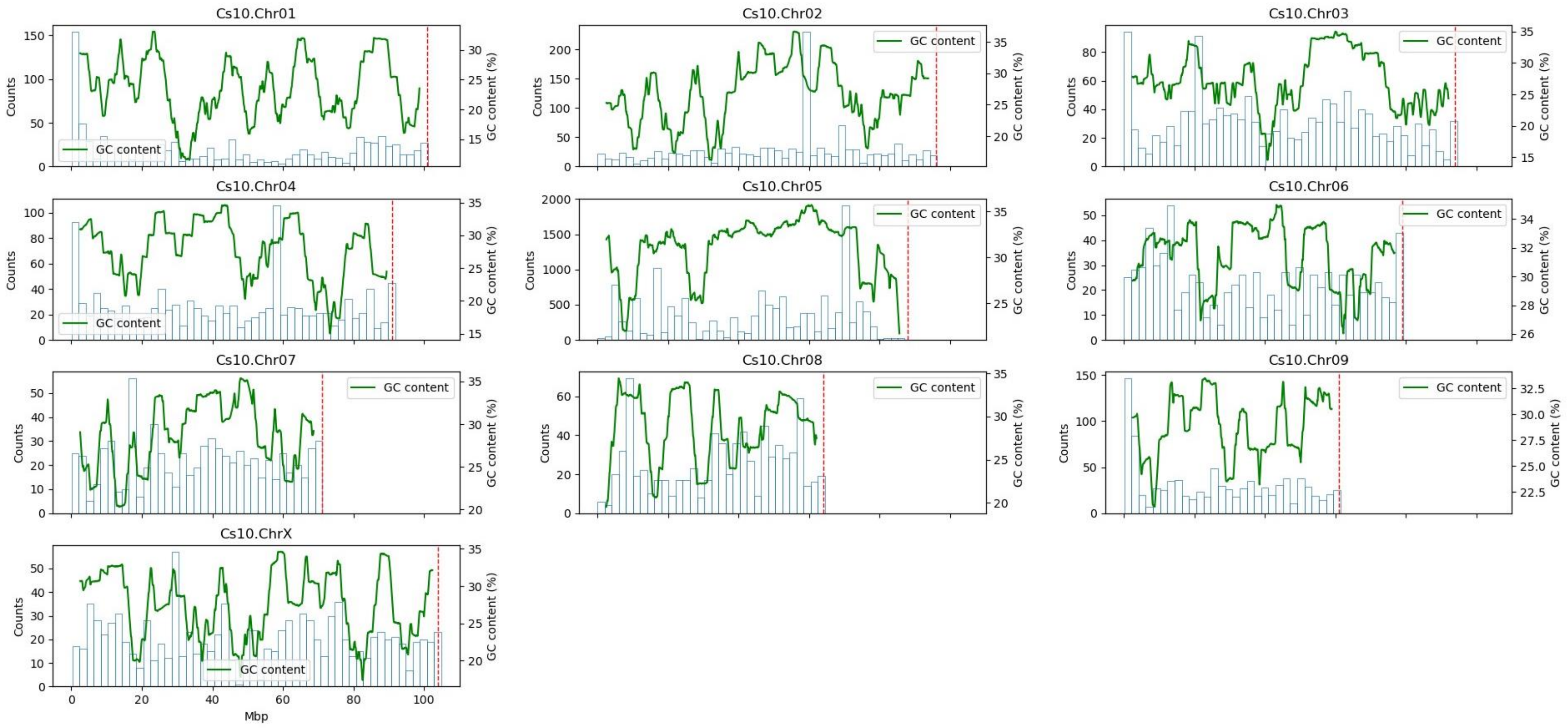


**Supplementary Figure 4**. GC content (green line) in 5 Mbp sliding window along ten cannabis chromosomes and counts of canonical mutants (bars) in 2 Mbp bins.

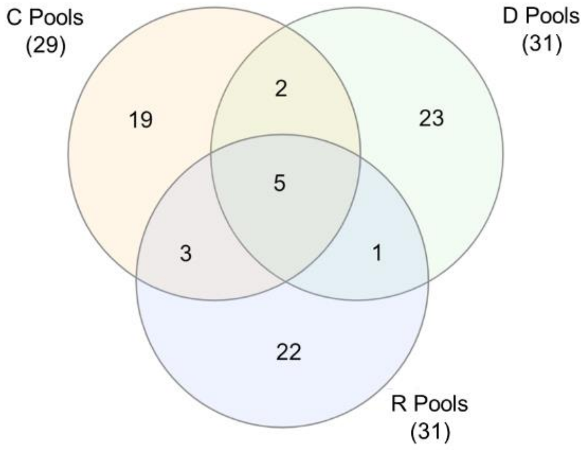


**Supplementary Figure 5.** Venn diagram of the candidate mutations identified in each pooling dimension after the application of the alternative allele frequency filter

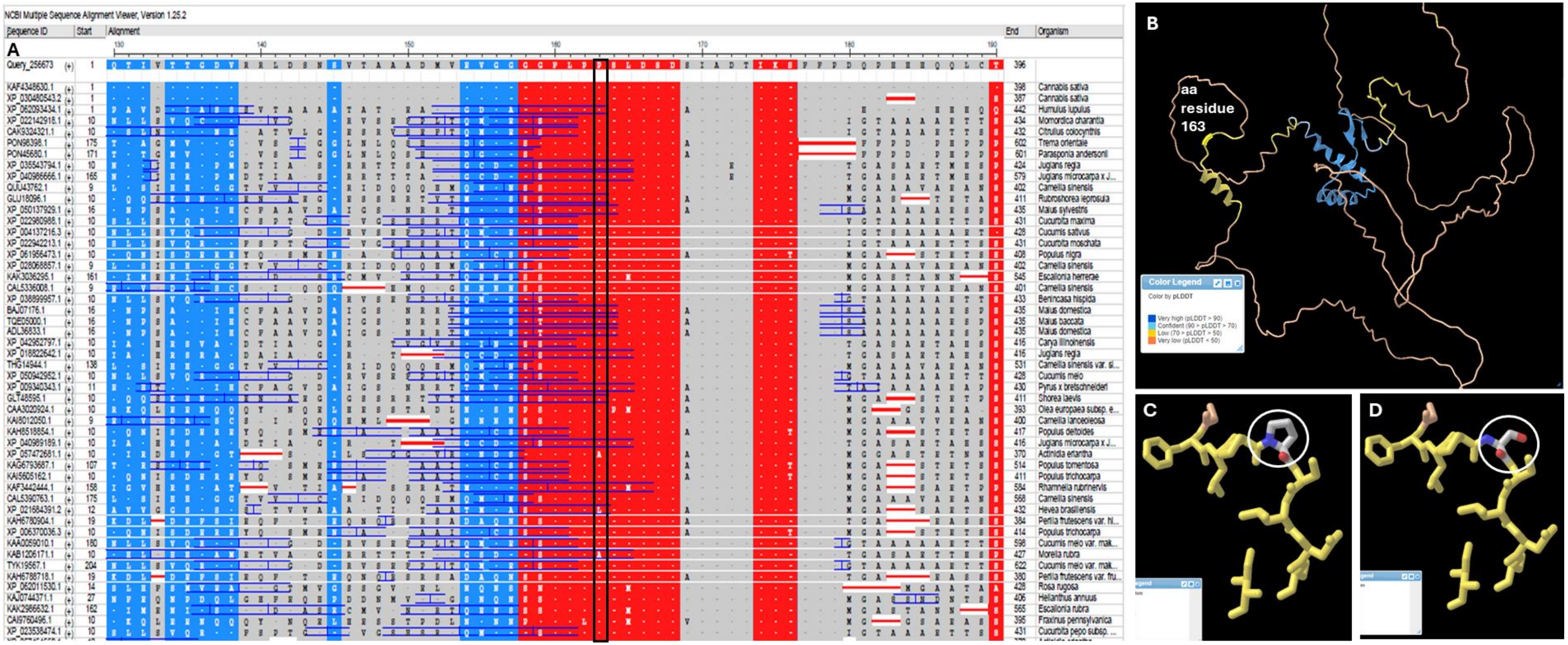


**Supplementary Figure 6**. Multiple alignment (A) using *CsTCP4* as anchor to compare sequence conservation from amino acid positions 130 – 190 across homologues in other species. Black box indicates the site of the mutation producing a P ->S substitution at position 163. Red indicates areas of high sequence conservation while blue and grey indicate less conserved regions. Protein structure as predicted by AlphaFold with position 163 in yellow (B) and effects on folding structure of mutation resulting in a P ->S substitution at position 163 (circled) in WT (C) and mutant (D) proteins.
